# Genomics epidemiology analysis reveals hidden signatures of drug resistance in *Mycobacterium tuberculosis*

**DOI:** 10.1101/2022.03.15.484552

**Authors:** P.M. Mejía-Ponce, E.J. Ramos-González, A.A. Ramos-García, E.E. Lara-Ramírez, A.R. Soriano-Herrera, M.F. Medellín-Luna, F. Valdez-Salazar, C.Y. Castro-Garay, J. Núñez-Contreras, M. De Donato-Capote, A. Sharma, J.E. Castañeda-Delgado, R. Zenteno-Cuevas, J.A. Enciso-Moreno, C. Licona-Cassani

## Abstract

*Mycobacterium tuberculosis* (*Mtb*) causes the majority of reported cases of human tuberculosis (TB), one of the deadliest infectious diseases worldwide. New diagnostic tools and approaches to detect drug-resistance must be introduced by 2025 to achieve the End-TB Strategy goals set for 2030 by the WHO. Genomic epidemiology of TB has allowed the expansion of catalogs listing genetic signatures of Mtb drug-resistance. However, very few *Mtb* strains from Latin America have participate in previous genomic epidemiologic efforts. Here we present the first functional genomic epidemiology study of drug-resistant *Mtb* strains in Mexico, incorporating the genomic characterization of 133 genomes, including 53 newly sequenced isolates, to provide a comprehensive phylogeographic analysis of drug resistant *Mtb* in Mexico. The study evidences the prevalence of Euro-American Lineage L4 (96.2%), featuring a uniform distribution of the sublineages X-type (33.08%), LAM (22.56%), and Haarlem (21.05%). Our results demonstrate low levels of agreement with traditional drug sensitivity tests (DST), raising concerns for drug-resistant isolates lacking any previously reported genetic signatures of resistance. Finally, we propose a novel functional networking tool (FuN-TB) to explore metabolic and cellular signatures of drug resistance. Applying functional genomics approaches to Latin American *Mtb* genomes will provide new drug-resistance screening targets that can contribute to bed side decision-making and advise local public policy.

**Abstract importance:** We presented the first phylogeographic analysis of *Mycobacterium tuberculosis* (Mtb) of Mexico. Our analysis integrates 133 genome sequences and is focused on the identification of genetic signatures associated to drug-resistance. The results show the geographic distribution of sublineages and drug-resistance phenotypic classes. Additionally, we propose a novel functional networking tool (FuN-TB) to explore metabolic and cellular signatures of drug resistance associated. We show for the first time that Mtb isolates from Mexico encode for region-specific genetic signarures of antimicrobial resistance. Applying functional genomics approaches to Latin American Mtb genomes will provide new drug-resistance screening targets that can contribute to bed side decision-making and advise local public policy.

## Introduction

Tuberculosis (TB) is a bacterial infectious disease caused by members of the *Mycobacterium tuberculosis* complex (MTBC) (1). After COVID-19, TB is the deadliest global pandemic, with 1.5 million casualties in 2020, including 214,000 people co-infected with the Human Immunodeficiency Virus (HIV) (2). Drug-resistant *Mycobacterium tuberculosis* (*Mtb*) is a concerning health problem given that 25% of annual TB-related deaths are linked to this cause (3). Globally, 3-4% of TB cases are resistant to either rifampicin or isoniazid, two of the most effective anti-TB drugs (2). In order to achieve the goals of its End-TB Strategy, the World Health Organization (WHO) has recently determined that new tools must be tested and introduced by 2025, including rapid point-of-care tests for detection and diagnosis, as well as global surveillance of drug-resistance (2). Molecular tools and bioinformatics with machine learning algorithms can also help deciphering new strategies for the epidemiological control of tuberculosis worldwide.

Designation of the drug-resistance markers in *Mtb* strains has derived from extensive genome-wide associated studies (GWAS) and genomic epidemiological projects performed in geographic regions in which TB is highly prevalent (4, 5). Recently, a list of 90 “canonical variants” associated with drug-resistant *Mtb* was published by the WHO, which was derived from the analysis of 38,000 MTBC genomes (6). In addition to complementing and increasing the resolution of drug-resistance genotyping, whole genome sequencing (WGS) studies also provide detailed information on sublineages and transmission groups of MTBC strains (7).

WGS-based technologies have transformed the field of TB epidemiology. Indeed, WGS is now considered the gold standard method for genotyping *Mtb* clinical strains in several countries (8, 9). However, in developing countries and regions with a lower incidence of TB, traditional genotyping techniques are still in use for diagnostics and drug-resistance detection. While expected, this situation demonstrates the need for increased genomic surveillance in order to understand the lineage distribution and evolutionary mechanisms that drive drug resistance at global scale. In addition to sequencing, it is also important to incorporate novel approaches during variant analysis to explore beyond the canonical gene markers associated with drug-resistance.

Although Mexico is considered a low-burden TB country, it ranks third in the top contributors of TB in the Americas (10). Most of the genotyping studies of *Mtb* clinical strains in this country were conducted using traditional techniques (11–17). To date, only two WGS-based studies of *Mtb* clinical strains from Mexico have been published (18, 19). In this study, we present a genotypic characterization of 53 newly sequenced clinical strains of *Mtb* from 17 representative states of Mexico. In addition to providing a robust identification of sublineages and drug-resistance patterns, we present the first phylogeographic analysis to integrate a total of 133 genomes, focused on drug-resistance phenotypes. Furthermore, we propose the use of a novel functional genomics comparative tool (FuN-TB) in order to identify unique SNP drug resistance-associated signatures and metabolic pathways, potentially related to pre-resistance phenotypes. While the number of genomes analyzed remains limited, our results highlight the importance of exploring the genomic diversity of Mexican and Latin American MTBC genomes in order to implement more effective drug-resistance control strategies at the global scale.

## Materials and methods

### Mycobacterial strains and culture

A total of 56 clinical samples of *Mtb* were obtained from sputum. These were collected between 1998 and 2019 from TB positive patients from 17 Mexican states: Nuevo León (n=13), Durango (n=7), Jalisco (n=6), Baja California (n=6), Ciudad de México (n=5), Zacatecas (n=4), Sinaloa (n=2), Oaxaca (n=2), Estado de México (n=2), Chiapas (n=2), Sonora (n=1), Puebla (n=1), Veracruz (n=1), Yucatán (n=1), Nayarit (n=1), San Luis Potosí (n=1), and Campeche (n=1). Transportation and processing of samples was conducted in adherence with the biosafety recommendations established by the Institute of Diagnosis and Epidemiological Reference (InDRE, by its Spanish acronym). The samples were isolated in Lowestein-Jensen medium (for two months at 37 °C) for phenotypic characterization and maintenance. Isolation and primary characterization were performed at the Biomedical Research Unit of Zacatecas (UIBMZ-IMSS).

### Phenotypic drug sensitivity test (DST) and in-silico spoligotyping analysis

A first-line antibiotic DST was performed with the fluorometric method BACTEC MGIT 960 (Becton-Dickinson, USA), using the following critical concentrations: isoniazid (H) = 0.1μg/mL; rifampicin (R) = 1.0 μg/mL; ethambutol (E) = 5.0 μg/mL; and streptomycin (S) = 1.0 μg/mL. A pyrazinamide (Z) test was performed using the BACTEC MGIT 960 PZA kit (Becton Dickinson). Second-line antibiotics DSTs were not performed due to infrastructure limitations. *In-silico* spoligotyping was performed with the SpoTyping tool (20).

### Genomic DNA extraction

The genomic DNA of *Mtb* colonies was extracted as previously described (21), with minor modifications. Briefly, the biomass from two-months growth in Lowenstein-Jensen tubes (BD, BBL) was resuspended in 400 μL of 1X TE buffer. Cells were inactivated by incubating the cell suspension at 80 °C for 20 min, followed by an overnight incubation at room temperature with 50μL of 10mg/mL lysozyme solution (Millipore-Sigma, USA). Cellular lysis was performed by adding 70μL of 10% SDS and 5μL of 10mg/mL Proteinase-K (Millipore-Sigma, USA), mixing and incubating at 65 °C for 10 min. Then, 100μL of 5M NaCl (Millipore-Sigma, USA) and 100μL of pre-warmed (65 °C) solution of CTAB/NaCl (40 mM/0.1 M) (Millipore-Sigma, USA) were added, and the suspension mixed vigorously, and incubated at 65 °C for 10 min. DNA extraction was achieved by adding an equal volume of chloroform-isoamyl alcohol (24:1) (Millipore-Sigma, USA) to the lysate, which was then mixed and centrifugated at room temperature. The aqueous phase was treated with 0.6 volumes of isopropanol (Millipore-Sigma, USA) to precipitate the DNA. The resulting DNA pellet was then washed twice with 500μL of cold 70% ethanol solution. After letting the pellet air dry for 10 min, the DNA was resuspended in sterile DNase-free water. DNA integrity was verified by electrophoresis in a 2% agarose gel (Millipore-Sigma, USA). DNA concentration and purity were determined by spectrometry with a Nanodrop 2000 (ThermoScientific, USA) and by fluorometry with a Qubit v.3 (Invitrogen, C.A., USA).

### Whole genome sequencing

Genomic libraries were prepared from 1 ng of high-quality genomic DNA using the Nextera DNA Flex Library Prep kit (now called Illumina DNA Prep, Illumina C.A., USA), following the manufacturer instructions. Library quality control was performed using the DNA 1000 kit (Agilent Technologies, USA) on a Bioanalyzer 2100 (Agilent Technologies, USA). Samples that passed the quality-control check were pooled and normalized according to the recommendations of the protocol. Prior to loading the cartridge, the pool was heated at 95 °C for 2 min to enhance the sequencing performance of the genomic regions rich in GC-content. After pre-heating, the pool was Tecnologico de Monterrey, Queretaro Campus, Mexico, using a NextSeq 550 (lllumina, C.A., USA) in a 2×150 paired end format.

### Quality control for the construction of a high-quality genome database of Mexican Mycobacterium tuberculosis isolates

In addition to the newly sequenced samples, in this study, we used a set of 80 high-quality whole genome sequences of *Mtb* previously reported in Madrazo-Moya et al. (18), with accession number PRJEB30933. For these samples, drug-resistance was tested by the fluorometric method (BACTEC, MGIT 960 Becton-Dickinson). *Mycobacterium bovis* was selected to root the phylogenetic trees and thus reads, corresponding to sample SRA: ERR2659159, were included in our analysis.

Quality control of the raw reads was examined using the software FastQC v0.11.9 (22) before and after applying Trimmomatic v0.33 (SLIDINGWINDOW:5:20) (23). Reads shorter than 20bp were excluded for the subsequent analysis. MTBseq pipeline v.1.0.3 (24) was used to obtain a list of high-quality variants from all samples, as well as the lineage classification, determination of transmission groups, and a concatenated list of SNPs for further phylogeographic reconstruction. We used MTBseq with the default parameters.

### Phylogeographic analysis

The phylogenetic tree was reconstructed using a concatenated SNPs list obtained from the MTBseq pipeline. The best substitution model was determined by MEGA X (25). RaxML-NG platform (26) was used to obtain the best phylogenetic tree, with a bootstrap cutoff of 0.03. Edition of the phylogenetic tree was conducted with the software iTOL v6 (27). The genotype results were visualized and integrated into the phylogenetic tree image using the iTOL-adapted files obtained from the TB-profiler analysis. Decimal geographic coordinates were determined for each of the Mexican states using the web portal https://www.geodatos.net. Phylogeography was obtained using a metadata file with all genomes together and the corresponding phylogenetic tree in the Microreact platform (28).

### Prediction of drug-resistant phenotypes

The drug-resistance genotype for both first-and second-line antibiotics was determined using the tool TB-profiler v4.0.3 (29) from the high-quality trimmed reads. Sensitivity, specificity, and positive (PPV) and negative (NPV) predictive values were determined with the “epiR” package from R. The strength of agreement between phenotype and genotype predictions of drug resistance was determined with the Cohen kappa (k) coefficient by using the “fmsb” package from R. A significance level of α=0.05 was used for all statistical tests. Functional enrichment analysis was performed with the DAVID Bioinformatics Resources 6.8 server (30), using the locus tag of the *Mtb* H37Rv genes of interest as input identifiers, and the default parameters.

### Genome-scale functional analysis of non-canonical variants associated with antibiotic resistance

Functional analysis of the genomic variants present in samples with specific phenotypic or epidemiological attributes was conducted using the FuN-TB (Functional Networks for TB genomes) tool. As input, this bioinformatic tool uses a list of high-quality variants, for example, those derived from the MTBseq pipeline, along with a list of sample-associated metadata or attributes (*e.g.,* phenotypic resistance, comorbidity, etc.). After selecting the attribute of interest, non-synonymous SNPs associated with functionally annotated genes are compared among samples classified by attribute. To identify the genes with the highest number of frequent mutations among the compared samples, the FuN-TB tool calculates a *fitness score* (ν) for each gene, defined as

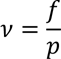

where *p* is the total number of SNPs within the gene and *f* is the sum of the frequencies of all SNPs in the gene. The higher the value (ν) obtained for a given gene, the greater the number of frequent variants it contains. The FuN-TB tool provides a list of the shared and unique mutated genes among the attributes selected, which includes their *ν* value and functional annotation (*i.e*., protein function, protein family, pathways, biological process, and reactions) produced by the Rapid Annotation using Subsystem Technology (31) and the Uniprot (32) servers. The FuN-TB tool output is a functional network (a file compatible with network visualization software, *e.g.,* Cytoscape [33]) designed to represent the shared and unique mutated genes among the selected attributes. For each network, the central nodes (selected attributes) are connected to peripheral nodes that represent the genes with SNPs. The functional network file also includes the *ν* values for each gene, which can be used as peripheral node traits and colored by gradient. FuN-TB tool algorithm is available in GitHub (https://github.com/ind-genomics).

### Molecular docking

Homology models of the mutated *GyrA* protein were constructed using the Swiss Model web server (https://swissmodel.expasy.org/), [34]). The 5BS8 crystallized protein from the Protein Data Bank (https://www.rcsb.org/) (35), which corresponds to the *GyrA* protein from *Mtb* H37Rv, was used as a template for model reconstruction. The UCSF Chimera program (v 1.12, [36]) was used to modify the amino acid sequences according to the mutations found in the DNA-seq experiments. Docking simulations were performed with AutoDock Vina (v. 1.1.2, [37]). The binding energy obtained for the first reference docking was used as a reference with which to infer the effects of mutated proteins on the ligand binding affinity. Structural analysis and plotting were conducted in the PyMOL software (v. 2.4.1, [38]).

### Ethical concerns

All of the information collected from TB patients was treated confidentially. Sampling was carried out without physical intervention, with full care given to the patient’s integrity. The study was approved by the National Scientific Research and Ethics Committee of the Instituto Mexicano del Seguro Social (IMSS), Mexico (R-2005-3301-18, and R-2013-785-001).

## Results

### Epidemiological, clinical and phenotypic characterization of newly sequenced Mtb samples from Mexico

We analyzed a total of 56 *Mtb* clinical isolates derived from patients with a positive diagnosis of pulmonary TB. The samples included represented 17 different Mexican states (Table 1). Overall, the mean age of the population sampled was 43.5 (range 16-85 years), with an approximately equal gender distribution (male: n=27, 48.2%). The most common comorbidity among the patients was Type-2 Diabetes Mellitus (T2DM) (n=11, 21%). Three isolates (6%) were from patients co-diagnosed with HIV. Of the total number of patients, 38% (n=20) had been vaccinated against TB, as verified by the presence of the characteristic scar of the Bacillus Calmette-Guérin (BCG) vaccine. Whereas 32% (n=18) of the patients were relapse cases of TB. **Supplementary Table 1** contains a full epidemiological description of the population sampled in this study.

**Table 1.** Epidemiological data of the *Mycobacterium tuberculosis* clinical samples (N=53). Type-2 Diabetes mellitus (T2DM); Not determined (ND); Human Immunodeficiency Virus (HIV); Latin American-Mediterranean (LAM).

| Characteristic | n (%) |
| --- | --- |
| **Sex** |  |
| Male | 27 (48.2) |
| Female | 18 (32.1) |
| ND | 11 (19.6) |
| **Age (years)** |  |
| 15-24 | 7 (12.5) |
| 25-54 | 26 (46.4) |
| >=55 | 12 (21.4) |
| ND | 11 (19.6) |
| **Comorbidity** |  |
| T2DM | 11 (19.6) |
| Alcoholism | 7 (12.5) |
| Smoking | 5 (8.9) |
| HIV | 3 (5.4) |
| **Treatment history** |  |
| Previously treated | 18 (32.1) |
| Newly diagnosed | 15 (26.8) |
| Unknown | 21 (37.5) |
| **TB vaccination** |  |
| BCG scar | 20 (35.7) |
| Unknown | 28 (50.0) |
| **Location** |  |
| Nuevo León | 13 (23.2) |
| Durango | 7 (12.5) |
| Baja California | 6 (10.7) |
| Jalisco | 6 (10.7) |
| Ciudad de México | 5 (8.9) |
| Zacatecas | 4 (7.1) |
| Chiapas | 2 (3.6) |
| Estado de México | 2 (3.6) |
| Oaxaca | 2 (3.6) |
| Sinaloa | 2 (3.6) |
| Campeche | 1 (1.8) |
| Nayarit | 1 (1.8) |
| Puebla | 1 (1.8) |
| San Luis Potosí | 1 (1.8) |
| Sonora | 1 (1.8) |
| Veracruz | 1 (1.8) |
| Yucatán | 1 (1.8) |

According to the DST, 31 samples (55.4%) were found to be “Susceptible” to all of the antibiotics tested. The remaining 23 samples showed a phenotype of drug-resistance against H (n=16), R (n=12), E (n=7), S (n=12), and Z (n=10). According to clinical sub-classification, eight samples (15%) were classified as “pre-MDR”, while ten samples (19%) were classified as multidrug-resistant (“MDR”). Five samples (9%) showed resistance to E, S, or Z (classified as “Other”). No conclusive results for drug resistance were observed for sample IMSS011, as well as specific pyrazinamide test results for samples IMSS001, IMSS002, IMSS018, and IMSS023.

### Lineage classification and phylogenetic analysis of newly sequenced Mtb samples from Mexico

From the initial collection of 56 *Mtb* clinical samples considered for sequencing, 53 had good-quality complete genomes. During library preparations, we were unable to obtain the genomic amplification for sample IMSS005. Furthermore, samples IMSS011 and IMSS036 were excluded from the analysis due to their low number of bases (110,090 bp and 2,185,476 bp, respectively) mapped against the reference genome *Mtb* H37Rv (NC_000962.3). For the remaining 53 *Mtb* genomes, an average genomic coverage of 104.5 (range: 303.3-to 20.97-fold) was obtained with respect to the reference genome. All sequencing-related data of the 53 *Mtb* genomes are presented in **Supplementary Table 2**.

The lineage classification showed that 98% of the samples belonged to the Euro-American Lineage (L4) while only one sample, collected from Sinaloa state, was classified as the Beijing Lineage (L2). The sublineages identified within our samples were X-type (n=18, 34%), LAM (n=12, 23%), mainly-T (n=10, 19%), Haarlem (n=8, 15%), Euro-American (n=2, 4%), S-type (n=2, 4%), and Beijing (n=1, 2%). In order to compare our results with traditional lineage determination techniques, we performed an *in silico* spoligotyping analysis. The results showed the differences expected, since the spoligotyping analysis indicated that, of the samples, 75% belonged to the Euro-American lineage, while 17% were Indo-Oceanic, 6% were East-Asian (Beijing) and 2% were *Mycobacterium bovis.* A comparison between the SNP-based analysis and the *in silico* spoligotyping for each sample is presented in **Supplementary Table 3.**

We performed a phylogenetic reconstruction for the 53 *Mtb* genomes using a total of 8,002 unambiguous SNPs, excluding variants from repetitive regions and drug-resistance markers (Fig. 1). The phylogeny structure showed a correct distribution of the samples according to their sublineage classification. Most of the samples classified as MDR belonged to the Haarlem sublineage. In addition, the X-type sublineage was prominent for presenting a greater number of samples showing at least one drug-resistance phenotype, as well as a higher number of samples derived from diabetic patients. On the other hand, the four samples with the HRESZ-resistance profile (IMSS009, IMSS040, IMSS041, and IMSS052), showed no correlation with respect to their sublineage classification. Considering an SNP distance of 12 bp, we found no transmission groups within this dataset.

**Figure 1.**
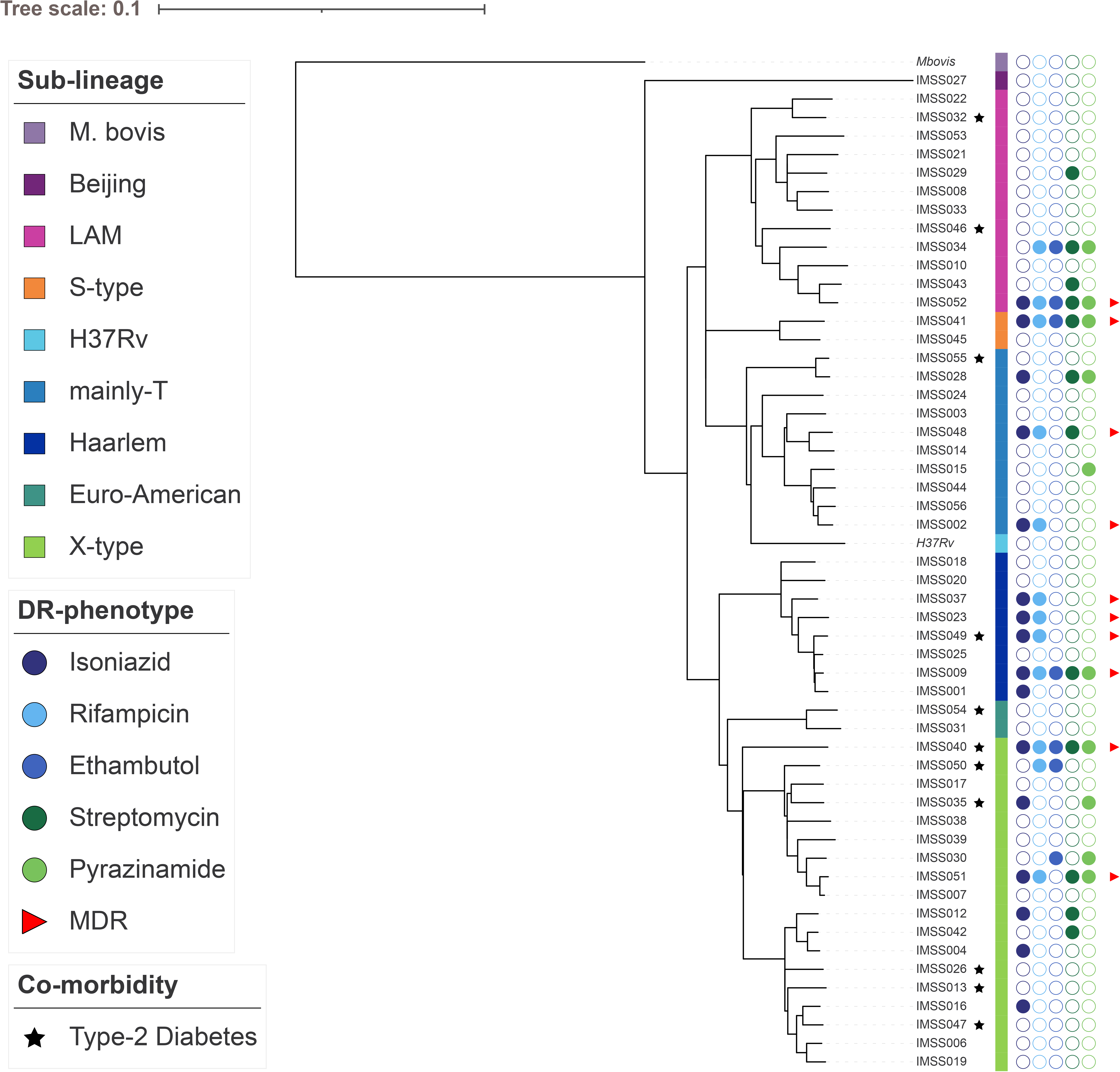
Phylogenetic reconstruction of the 53 clinical samples of *Mtb* from Mexico. A total of 8002 SNPs were used for this phylogeny, which were filtered and concatenated by MTBseq pipeline. The best substitution model was GTR+FO+G4m. Considering a cutoff of 0.03, tree bootstrapping converged after 50 replicates.

### Identification of first- and second-line multidrug resistance associated SNPs

For readability, we refer to all drug-resistance-associated variants (both first- and second-line antibiotics) identified by TB-profiler tool as “canonical variants”. A total of 50 such canonical variants were identified in 19 of the 53 genomes (Table 2). The most frequent variants associated with H resistance were *ahpC_*Asp73His, *fabG1_*–15C>T, and *katG_*Ser315Thr, while *rpoB_*Asp435Val, *rpoB_* His445Asn, and *rpoB_* Leu452Pro were the most frequent mutations related to R resistance. For E resistance, the mutations *embB_*Met306Ile/Leu/Val and *embB_*Gly406Ala were identified. In case of S resistance, only three variants were found: *gid_*Leu79Ser, *rpsL_*Lys43Arg, and *rrs_*514a>c (RNA level). For Z resistance, six samples showed different variants within the *pncA* gene, including two insertions (391-392insGG and 118-119insGG), one deletion (2288707-2291359 at DNA level) and three SNPs (Thr61Pro, Trp68*[stop codon], and Gly132Ser).

**Table 2.**
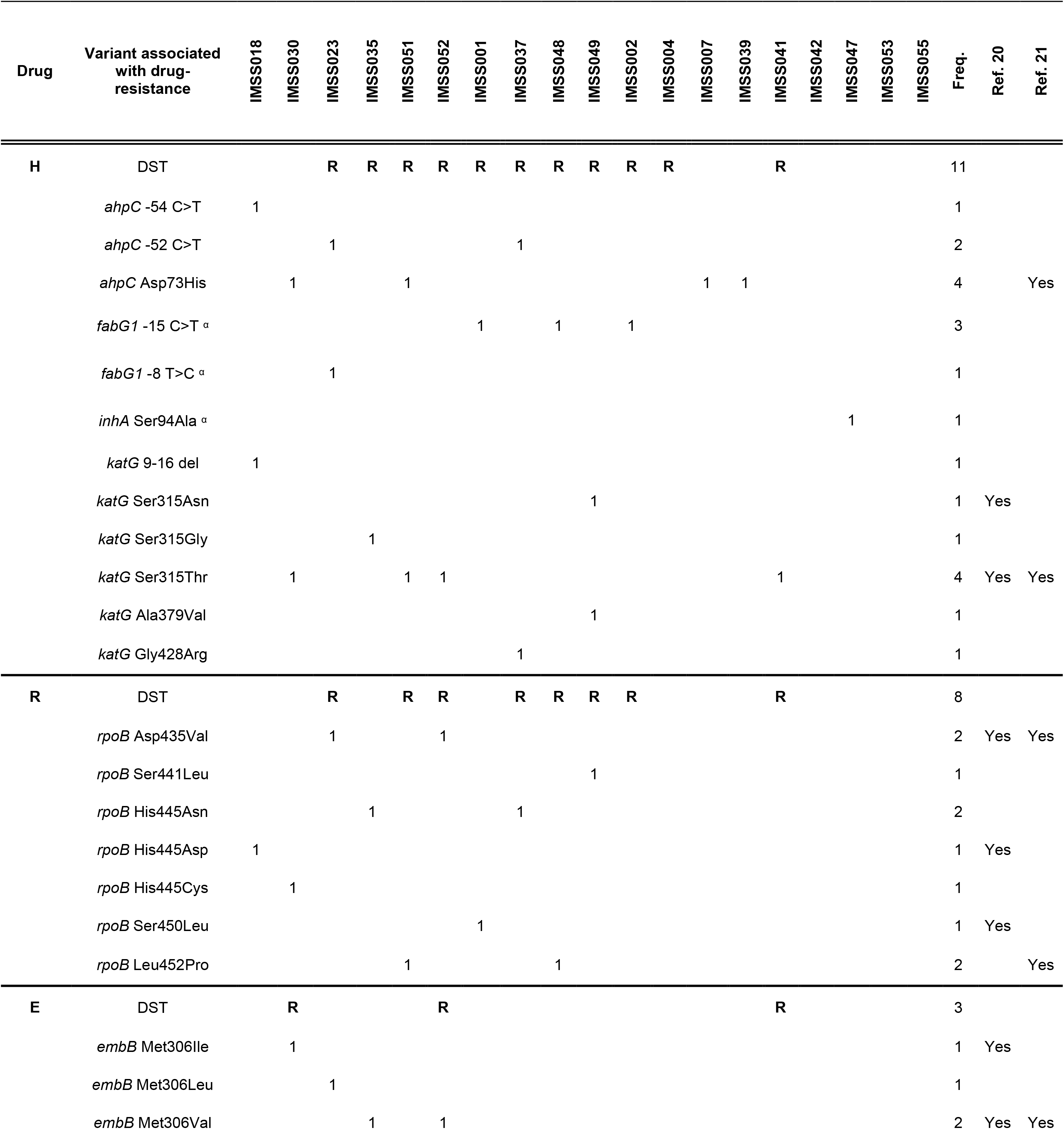
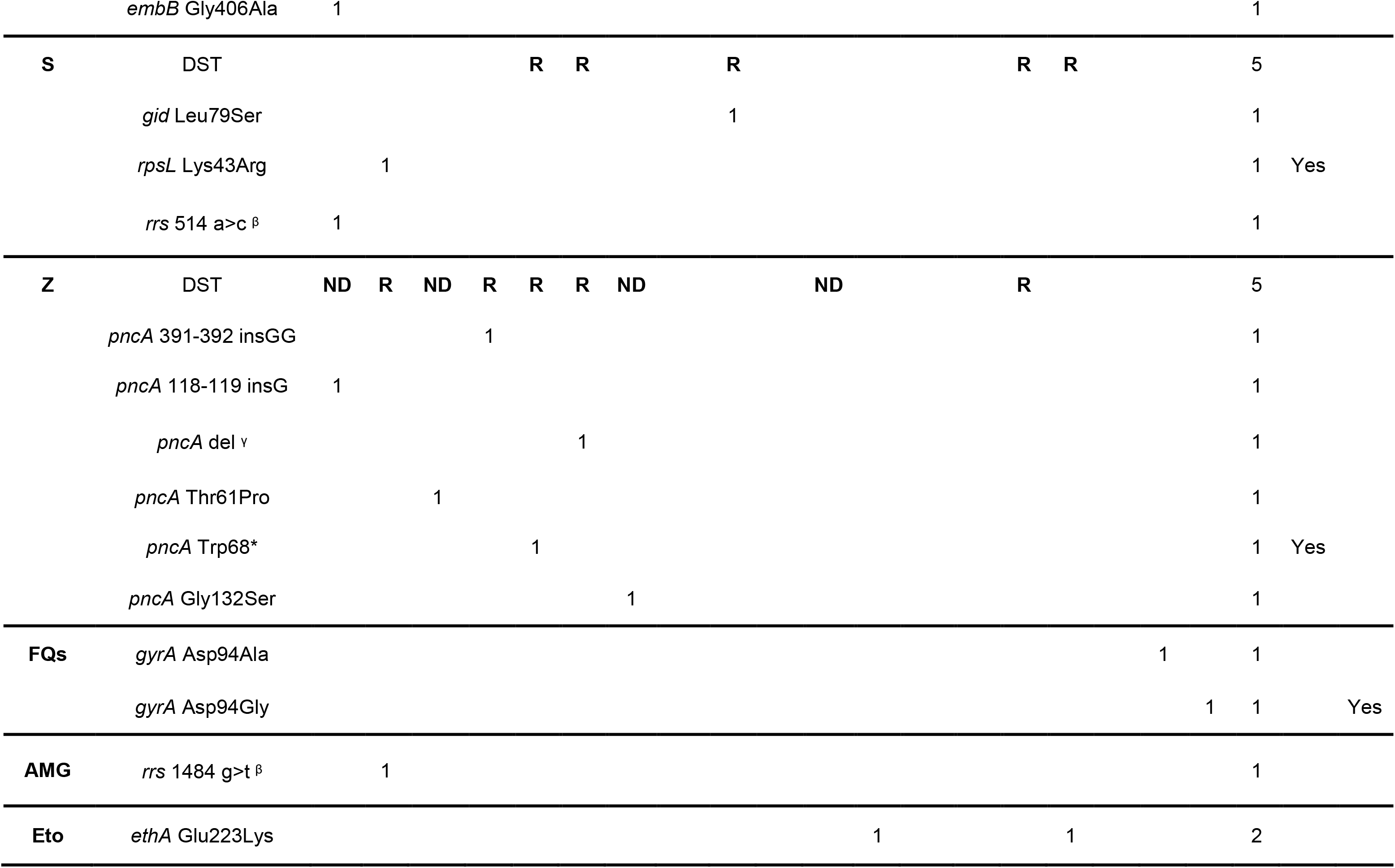
Canonical variants associated with drug-resistance in the 53 clinical samples of *Mtb*. α Mutations associated with ethionamide resistance, also; β Mutations at RNA level; γ Deletion at chromosome level from 2288707 to 2291359 positions; *Stop codon; DST = Drug Sensitivity Test, phenotypic assay based on the MGIT960 technique.

In five samples, we identified second-line antibiotic-associated variants: *gyrA_*Asp94Ala/Gly, *rrs_*1484g>t (RNA level), and *ethA_*Glu223Lys, which are associated with resistance to fluoroquinolones, injectable aminoglycosides (kanamycin, capreomycin, amikacin), and ethionamide, respectively. We performed a molecular docking simulation between the *GyrA* protein (antibiotic target) and the antibiotic moxifloxacin. We predicted the effect of the variants identified on the GyrA-moxifloxacin complex and found that some non-canonical variants modified the *GyrA* topological polar surface area (TPSA), reduced the predicted binding energy values, and generated bulky obstacles within the binding site. The results of the molecular docking simulation from the GyrA-moxifloxacin complex are presented in **Supplementary Fig. 1**.

### Discrepancies between phenotypic and genotypic drug-resistance predictions in *Mtb* strains from Mexico

The correlation between the phenotypic and genotypic predictions of drug-resistance for our *Mtb* isolates, showed a sensitivity and specificity for H of 67% (C.I. 38-88%) and 84% (C.I. 69-94%), respectively. For R, were 60%, (C.I. 26-88%) and 86%, (C.I. 72-95%), respectively. While specificity for E and S were 90% (C.I. 77-97%) and 78% (C.I. 64-88%), respectively), sensitivity values were 40% (C.I. 5-85%) and 33% (C.I. 1-91%), respectively. Four strains (IMSS001, IMSS002, IMSS018, and IMSS023) were omitted from this statistical analysis for Z since we did not have their corresponding phenotypic data, although most of these strains contained canonical variants of Z resistance. Taking these factors into account, the sensitivity and specificity values for Z were 100% (C.I. 29-100%) and 85% (C.I. 71-94%), respectively.

In general, the positive predicted values (PPV) were poor for all antibiotics tested (mean=35.8%, range=8-62%), whereas the negative predicted values (NPV) were relatively high (mean=92.8%, range=86-100%). The strength of agreement for H, R, and Z was defined as ‘Moderate’, as reflected by their k coefficient value (average k=0.45). In the case of E, the strength of agreement was determined as ‘Fair’ (k=0.25, C.I. −0.23, 0.73). The lowest ranked k value was observed for S (k=0.05, C.I. −0.40, 0.5), which indicated a strength of agreement defined as ‘Slight’. Table 3 presents the complete statistical data for the 53 *Mtb* samples.

**Table 3.**
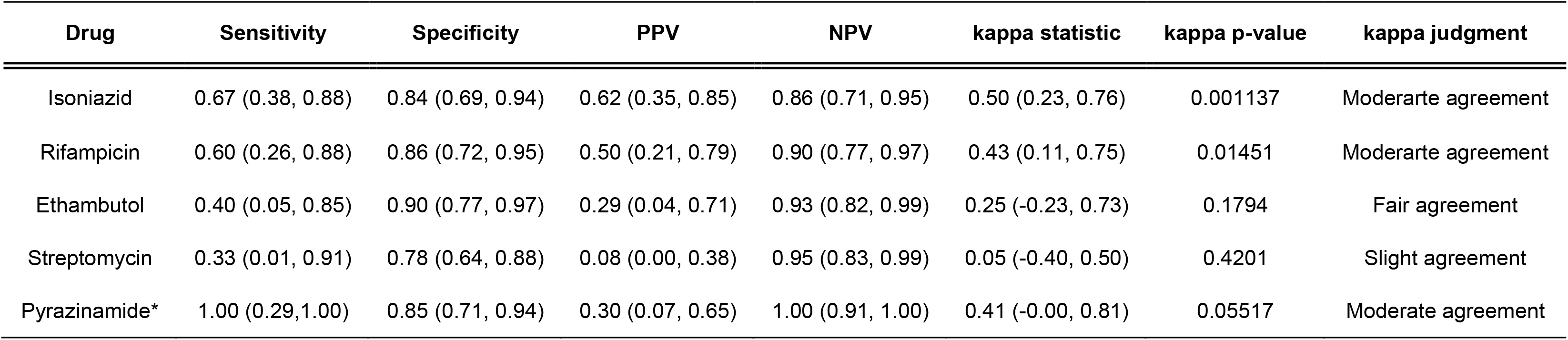
Statistic metrics related to the concordance between the phenotype and genotype of the 53 clinical samples of *Mtb*. The 95% confidence interval is indicated in parentheses. (*) Four samples were omitted from this analysis due to lack of phenotypic data.

Moreover, we found that several samples showed phenotypic susceptibility to certain antibiotics, but harbored corresponding canonical variants. For instance, six samples (IMSS007, IMSS018, IMSS30, IMSS039, and IMSS047) exhibited susceptibility to H but contained the canonical variants *ahpC_*Asp73His, *ahpC_*-54c>t, *katG_*9-16del, *katG_*Ser315Thr, and *inhA_*Ser94Ala, respectively. Other samples (IMSS001, IMSS018, IMSS030, and IMSS035), showing susceptibility to R, harbored mutations within the *rpoB* drug-resistance gene (positions Ser450Leu or His445Asp/Cys/Asn). Two S-susceptible isolates (IMSS18 and IMSS30) encoded for canonical variants *rrs_*514a>c and *rpsL_*Lys43Arg. In addition, three E-susceptible samples (IMSS018, IMSS023, and IMSS035) contained variants in the gene *embB* (Gly406Ala and Met306Leu), whereas two Z-susceptible samples (IMSS018 and IMSS23) encoded *pncA* 118_119 insG and Thr61Pro mutations, respectively.

We also found 33 samples that presented drug-resistant phenotypes but lacked the corresponding canonical variants. This was the case for six samples showing H resistance, six samples with an R-resistant phenotype and 21 samples exhibiting S-, E-or Z-resistant phenotypes (n=11, n=5, and n=5, respectively). More importantly, samples IMSS034, IMSS041, IMSS009, and IMSS040, which presented a resistant phenotype against >4 antibiotics, lacked all of their associated canonical variants. We have summarized all these genotype-phenotype discrepancies in **Supplementary Table 4**.

### Phylogeographic analysis showed a wide distribution of drug-resistant *Mtb* phenotypes in Mexico

We performed a WGS-based phylogeographic analysis focused on the drug-resistance of 133 *Mtb* clinical strains from Mexico, including newly sequenced samples (n=53) along with another 80 high-quality genomes previously reported by Madrazo-Moya et al., (18). A phylogenetic tree of the 133 *Mtb* genomes was reconstructed from 11,166 concatenated SNPs, excluding variants associated with drug-resistance and within repetitive regions (Fig. 2.A). The structure of the phylogeny was correctly distributed according to the sublineage classification of the samples. We found that all sublineages identified in Mexico had at least one sample classified as MDR. Moreover, each sublineage, apart from the S-type, had at least two samples derived from patients diagnosed with T2DM (34%, n=45). Interestingly, the most common drug-resistance classes found among the diabetic-derived samples were MDR (n=16) and pre-MDR (n=13). We also expected that some of our samples would probably fit into the transmission clusters previously identified in Veracruz (18), but that was not the case.

**Figure 2.**
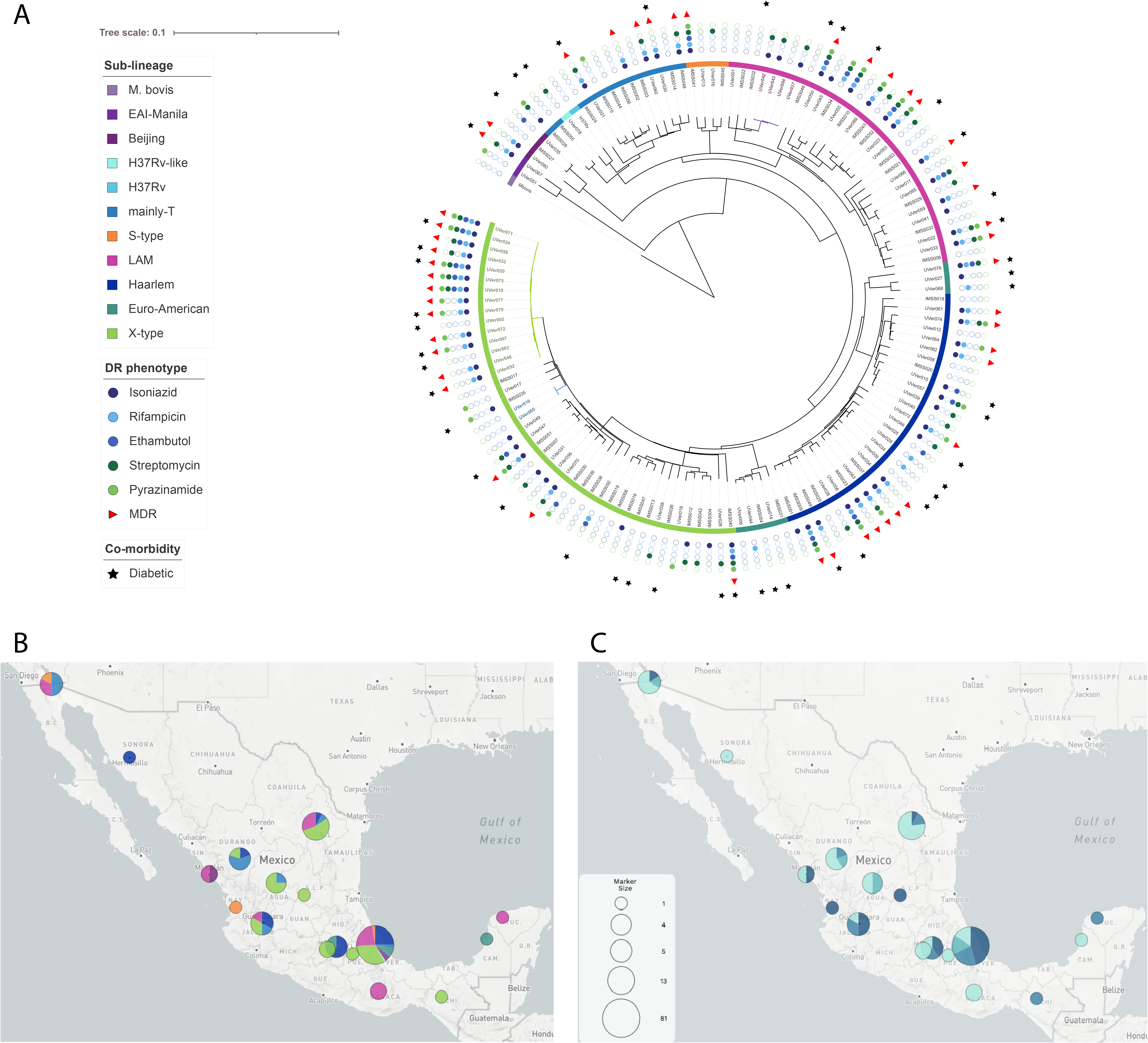
Phylogeographic analysis of 133 *Mtb* whole genomes from Mexico. A) Phylogenetic tree of 133 *Mtb* samples, including newly sequenced samples from this study and 80 *Mtb* genomes previously reported in (18). The phylogeny was reconstructed with 11,166 SNPs using the GTR+FO+G4m substitution model and a bootstrapping cutoff of 0.03, equivalent to 50 replicates. Sample labels colored in green, blue and purple indicate the three transmission groups previously identified in (18). B) Geographic distribution of the 133 *Mtb* genomes according to their sublineage classification. C) Geographic distribution of the 133 *Mtb* samples showing their drug-resistance classification.

In Figure 2.B we present the geographic distribution of the *Mtb* sublineages in Mexico. The main sublineage, X-type (33%), was found distributed among the states of Veracruz, Nuevo León, Zacatecas, Jalisco, Estado de México, Durango, San Luis Potosí, Puebla, and Chiapas. The LAM sublineage (23%) was identified in Veracruz, Nuevo León, Baja California, Oaxaca, Jalisco, Sinaloa, and Yucatán, while the Haarlem sublineage (21%) was located in Veracruz, Ciudad de México, Jalisco, Durango, Nuevo León, and Sonora. The less abundant lineages Beijing (L2, n=3) and EAI-Manila (L1, n=2) were only found in Veracruz and Sinaloa.

From this dataset of 133 *Mtb* samples, the most representative drug-resistance phenotypic clinical class was MDR (35%, n=47), followed by “Susceptible” (32%, n=43), “pre-MDR” (20%, n=26), and “Other” (13%, n=17). The geographic distribution of the phenotypic drug-resistance classes of the 133 *Mtb* samples from Mexico is presented in Figure 2.C. Samples classified as MDR and pre-MDR were mainly from the states of Veracruz, Jalisco, Nuevo León, and Ciudad de México. We found no correlation between sublineage and phenotypic drug-resistance class in this set of samples.

### Functional network analysis of non-canonical variants provides a broader metabolic and cellular scope of mono-resistant phenotypes

To understand the possible mechanisms of acquisition of, or adaptation to, a drug-resistant condition, we carried out a functional analysis of the non-canonical variants of the 133 *Mtb* genomes available from Mexico. An initial list of 24,272 variable positions was obtained from the variant calling analysis. A total of 5,230 positions classified as “uncovered” (Unc) by the MTBseq software were excluded from subsequent analysis since they belonged to low-quality sequence regions (*e.g*., repetitive regions). The resulting 19,042 variable positions were used as input to perform a functional network analysis with the FuN-TB tool (See **Methods**) in order to identify common mutated genes among samples that met certain attributes, as well as to prioritize the genetic function with their fitness score (*ν*). In this case, the attribute selected was a mono-resistant phenotypic profile. Thus, our working dataset comprised 28 samples that exhibited drug-resistance exclusively to H (n=8), R (n=4), E (n=2), S (n=10), or Z (n=4). To ensure that all of the variants we were going to analyze were associated with drug-resistance, we used the FuN-TB tool to filter the variants present in samples classified as “Susceptible”, both phenotypically and genotypically. A total of 3,307ble positions were obtained from this process, which served as the background for subsequent comparisons among the mono-resistant samples.

As a result of the comparison analysis of mono-resistant samples, a total of 743 non-synonymous SNPs belonging to 323 genes was obtained, and these are represented in a functional network presented in Figure 3. We found that the mutations *hpt_*Leu75Met, *etgB_*Pro193Gln, and *scoB_*Pro40Leu were convergent between different samples belonging to E, R, and S mono-resistant conditions. The gene *hpt* is involved in purine ribonucleoside salvage, whereas *etgB* is classified as an iron (II)-dependent oxidoreductase and *scoB* is related to the response to hypoxia. Similarly, mutations of *ligC_*Arg313His and *pe35*_Glu99* (*=stop codon) were convergent among samples belonging to E, H, and R mono-resistant conditions. The *ligC* gene is related to DNA repair and recombination, whereas *pe35* belongs to the PE-protein family. Interestingly, a list of 31 genes (*ν* range = 2.0 - 5.0) showed the same mutation between samples belonging to the E and R mono-resistant conditions. These mutated genes were involved in Actinobacterium-type cell wall biogenesis (*fadD19, fadD17, whiB3, ppsC,* and *ddlA*), ESX-n secretion system proteins (*espJ, eccE1,* and *eccA2*), the response to hypoxia (*ppk1*), transmembrane transport (*dctA*), chromosome segregation (*scpA*), biological processes involved in interaction with the host (*cyp125*), and amino acid biosynthetic processes (*proB* and *hsaF*), among others.

**Figure 3.**
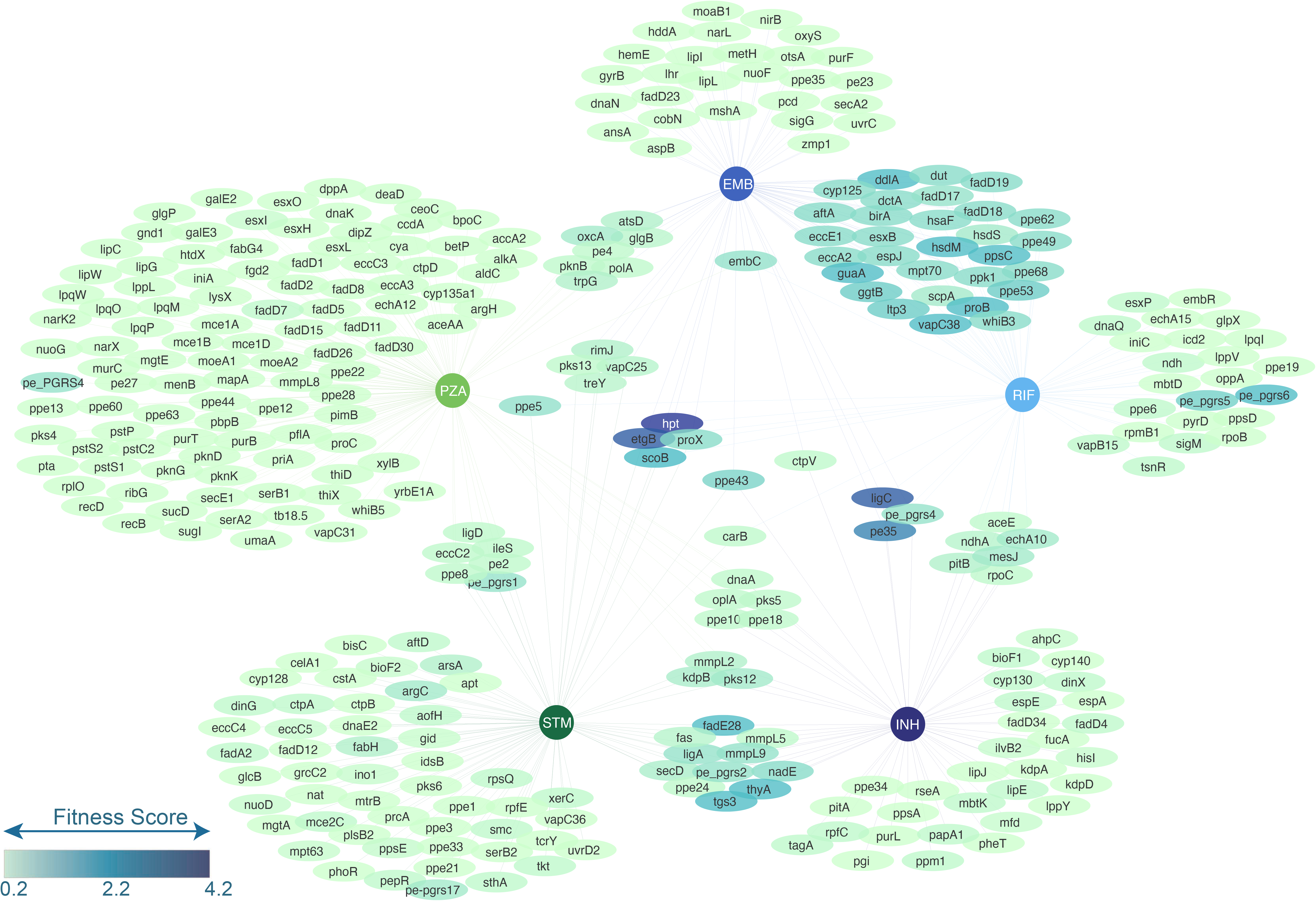
Functional network of mutated genes found in the mono-resistant dataset comparison. The five concentric nodes indicate the antibiotic to which each mono-resistant sample subset belongs. The color gradient of each gene indicates the fitness score (v) value. The higher v value for a gene, the greater number of frequent variants contains.

Besides the lack of correlation between the mode of action of H and S antibiotics, corresponding mono-resistant samples shared the same variable on genes *fadE28, tgs3, thyA, nade,* and *mmpL9*, which were involved in interaction with the host, response to hypoxia, response to antibiotics, NAD biosynthetic process, and evasion of host immune response, respectively. Genes *pks12*, *kdpB*, and *mmpL2* were found to be variable between H, S and Z mono-resistant phenotypes, but their low *ν* values indicated that they do not converge at the same variable positions within the gene. We found 41 mutated genes belonging to PPE, PE-PGRS, and PE family proteins (n=25, n=11 and n=5, respectively), which are commonly acknowledged as glutamine-rich proteins involved in interaction with the host.

Regarding the H-resistant condition, 32 unique genes appeared to be mutated with low ν values (*ν* range = 1.0 −2.0). Among these, 11 genes were expressed in, or related to, cell wall and plasma membrane processes (*lipE, ppm1, bioF1, cyp130, ahpC, kdpA, ppsA, lppY, pitA, ppe34*, and *pheT*), three genes were involved in DNA repair processes (*tagA, dinX*, and *mdf*), and two genes were related to amino acid biosynthesis (*ilvB2* and *hisI*). In the R-resistant condition, a total of 23 unique genes were found, including *sigM, tsnR, dnaQ, esxP, vapB15, ppe6, pe-pgrs6*, and *ndh* with low ν values (ν range = 1.0 −2.0). For the E-resistant condition, 27 unique genes were identified with very low ν values (ν=1.0). Of these, 14 genes are expressed, or involved, in cell wall processes. A total of 29 genes were found to be mutated exclusively in the S-resistant condition, including six genes involved in lipid metabolic processes (*fadH, ppsE, pks6, idsB, mgtA*, and *grcC2*). With respect to the Z-resistant condition, 106 unique genes were found to be mutated, with very low ν values (ν = 1.0). As in some of the previous cases, most of these genes (n=59) were involved in cell wall, plasma membrane, and extracellular region processes. The complete functional annotation of the 323 mutated genes, as well as their ν values, are presented in **Supplementary Table 5**.

### Functional network analysis of non-canonical variants in multidrug-resistant strains helps identification of pre-resistant metabolic signatures

To identify pre-MDR resistant cellular and metabolic signatures, we performed a functional network analysis with the FuN-TB tool, focusing on the non-canonical variants from samples that were mono-resistant to H (n=8) and R (n=4), as well as 11 MDR samples (resistant to H and R only). We used the same background of variants derived from the “Susceptible” samples obtained previously to exclusively analyze the variants associated with drug-resistance. As a result, a list of 520 non-synonymous SNPs corresponding to 299 genes was obtained. These are represented in a functional network in Figure 4.

**Figure 4.**
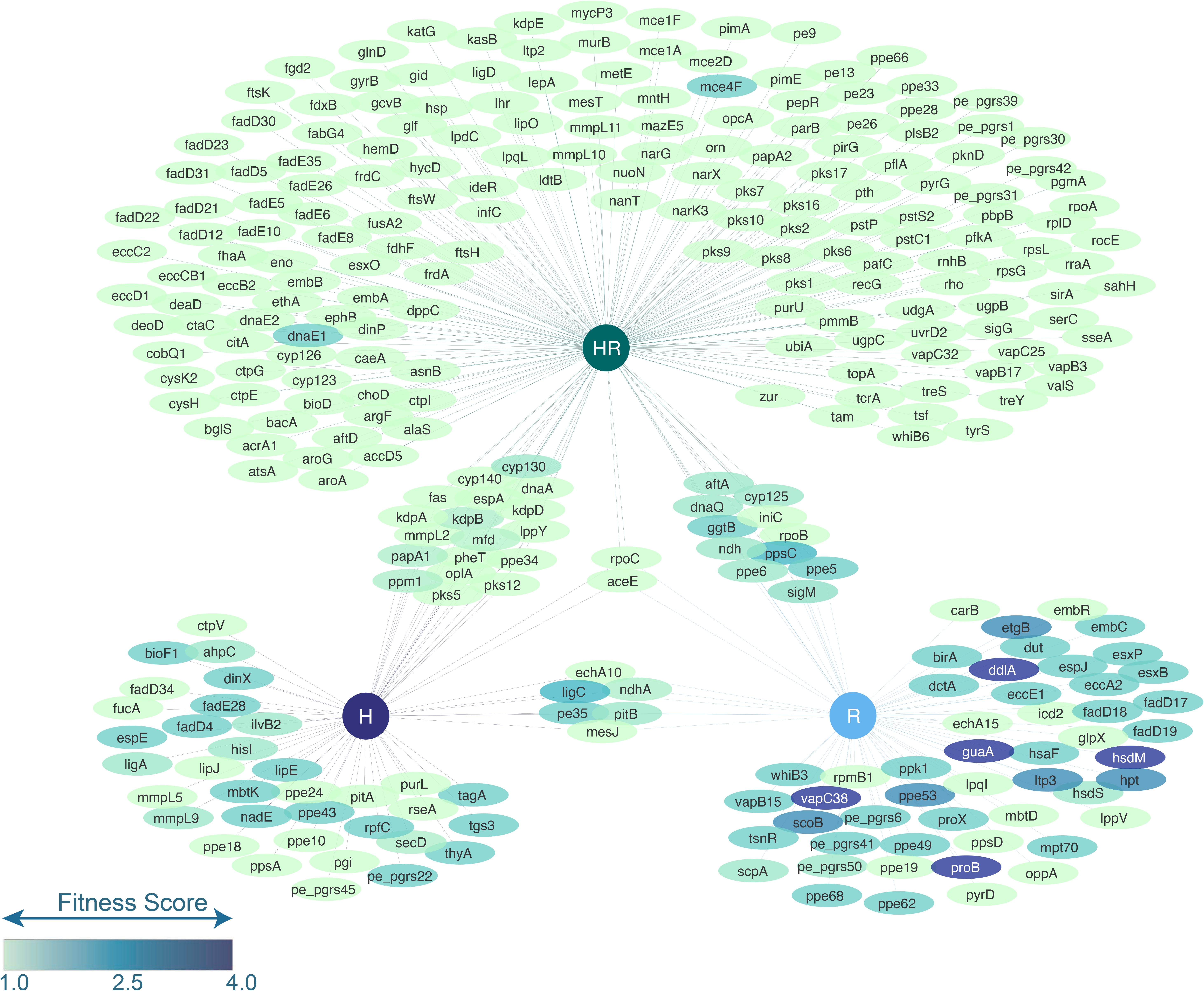
Functional network of the mutated genes found in the MDR dataset comparison. The HR central node corresponds to the sample dataset showing resistance against isoniazid and rifampicin, whereas H and R nodes indicate a corresponding mono-resistant phenotype. Color gradient of each gene indicates the fitness score (ν) value. The higher the ν value for a gene, the greater number of frequent variants it contains.

Functional network analysis identified *aceE* (part of the glycolytic process) and *rpoC* (member of the *Mycobacterium* virulence operon involved in DNA transcription) with mutations for the three subsets of samples (HR, H and R); however, we observed a low value (ν=1.0). Among the subsets HR and H subsets, a list of 18 shared mutated genes were found (range ν = 1.0 - 1.5), including two cytochrome P450 coding genes (*cyp130* and *cyp140*), two genes involved in potassium homeostasis (*kdpB* and *kdpA*), three genes related to fatty acid biosynthesis (*fas, pks12, pks5*), one transmembrane transport gene (*mmpL2*), and one gene participating in symbiont modulation of the host immune response (*ppe34*), among others. Except for *cyp130* and *pks5*, all of the genes mentioned above showed the same point mutations regardless of whether they belonged to the HR or the H subset. With respect to the samples from HR and R subsets, 11 mutated genes were found in common (range ν = 1.0 - 2.5). Some of these genes are involved in Actinobacterium-type cell wall biogenesis (*ppsC*), the response to xenobiotic stimulus (*sigM*), a cytochrome P450 encoding gene (*cyp125*), an isoniazid inducible encoding gene (*iniC*), and two members of the PPE family of proteins (*ppe5* and *ppe6*), among others. Comparison of these genes showed different variable positions between samples HR and R. Six mutated genes were shared between H and R mono-resistant samples (range ν = 1.0 - 2.5) and were related to DNA repair and recombination (*ligC*), phosphate ion transport (*pitB*), transmembrane transport (*ndhA*), tRNA modification or processing (*mesJ*), fatty acid metabolism cluster (*echA10*), and PE family members (*pe35*). From this group of mutated genes, only *ligC, pe35, echA10*, and *mesJ* retained the same point mutations between samples classified as R or HR resistant.

Approximately 60% of mutated genes (n=178) were only found in samples from the HR subset. The main significant enrichment clusters among these mutated genes were fatty acid biosynthetic processes (score=1.88, genes=8); secondary metabolite biosynthesis, transport, and catabolism (score=1.72, genes=6); metabolic process (score=1.54, genes=9); lipid homeostasis (score=1.49, genes=6); cell membrane (score=1.34, genes=36); DNA repair (score=1.19, genes=5); response to antibiotics (score=1.08, genes=7); ABC transporters (score=1.02, genes=10); cell wall organization (score=1.01, genes=7); cell division (score=0.99, genes=5); aminoacyl-tRNA biosynthesis (score=0.90, genes=3); and Glycolysis/Gluconeogenesis (score=0.68, genes=4). The complete functional annotation of the 299 mutated genes, as well as their ν values are available in **Supplementary Table 6**.

## Discussion

TB incidence in Mexico was estimated at 29,000 cases in 2018, of which 3.3% were reported as drug-resistant (39). The geographic distribution of TB cases in Mexico has been reported previously (40–45), but none of these studies provided genomic information about lineage and drug-resistance patterns of the isolates. Here, we presented a genotypic characterization of 53 newly sequenced clinical strains of *Mtb* from 17 representative states of Mexico. In addition to providing robust identification of sublineages and drug-resistance patterns, we presented the first phylogeographic analysis that integrates 133 genome sequences focused on drug-resistance phenotypes. Our phylogeographic analysis results show the geographic distribution of sublineages and drug-resistance phenotypic classes of 133 clinical samples of *Mtb*. The states of Veracruz and Nuevo León contributed the highest number of samples per site (n=81 and n=13, respectively), whereas Sonora, Nayarit, San Luis Potosí, Puebla, Chiapas, Campeche, and Yucatán provided only one sample per site. We also incorporated a functional network analysis of the non-canonical variants in order to identify metabolic and cellular signatures of resistance as a novel approach to the analysis of mutations that might otherwise be missed in global databases.

Regarding the sublineage classification of the 53 *Mtb* genomes, we found significant differences between the results obtained from the *in silico* spoligotyping and the SNP-based approach. The *in silico* spoligotyping indicated that most of the samples belonged to the Euro-American lineage (L4). Still, spoligotypes such as *M. bovis*, East-Asian (Beijing) and Indo-Oceanic were designated for 12 samples. Conversely, the SNP-based approach classified all but one sample (identified as Beijing) within the L4 lineage, including X-type (n=18) and LAM (n=11) sublineages as the most frequent.

Interestingly, we identified seven isolates (distributed in the north: Nuevo Leon, Zacatecas, San Luis Potosi; center: State of Mexico; and south: Chiapas) belonging to the X-type (4.1.1.3), an endemic sublineage recently reported in Veracruz and Guadalajara (18, 19) that has been strongly associated with drug-resistance (46). Three of these samples (43%) showed phenotypic drug-resistant profiles (EZ, HZ, and HRSZ, respectively). This data confirms the wide distribution of this sublineage in the country and the strong tendency to develop drug resistance. Overall, a comparison between the *in silico* spoligotyping and WGS indicated that the WGS-based approach increases the resolution for epidemiological typing of *Mtb* strains. These data contributed to a clearer understanding of the sublineages circulating in Mexico and offer the possibility for their inclusion within the global databases and use in comparative analysis.

The phenotypic results of drug-resistance against first-line antibiotics of the 53 *Mtb* samples indicated moderate resistance levels for H (30%), R (23%) and S (23%), but low resistance levels for E (13%) and Z (19%). In addition to the fact that 58% of samples were classified as phenotypically “Susceptible”, 19% were identified as MDR. Compared to 2013, national MDR rates were 11.1% (based on BACTEC assays), considering 18 Mexican states (47), which could suggest an increase in the MDR rates in the region. In Mexico, most drug-resistance phenotypic tests are conducted only for first-line antibiotics (13, 17, 40, 48). In addition to the growing prevalence of MDR strains circulating in the region, only one study has reported second-line antibiotics DST assays (49). Soon, it will be necessary to establish accessible methodologies, related to resources and infrastructure, at the main diagnostic points in order to identify and adequately report strains with extensively drug-resistance (XDR) profiles in Mexico.

With regard to drug-resistance genotypic analysis, we found some of the most frequent canonical variants, such as *ahpC*_Asp73His, *katG*_Ser315Thr, *rpoB*_Asp435Val and *rpoB*_Leu452Pro (18, 19). However, a total of 24 canonical variants reported here were shown in Mexico for the first time (Table 2). The WGS technique has been recognized to significantly increase the resolution of the phenotypic DST in *Mtb* samples (7). Unfortunately, and in contrast to some other countries, representation of *Mtb* genomes from Mexico is limited in global databases (6). The lack of genomic information limits accurate sublineage classification and detection of drug-resistance variants in the region. Mexico hosts one of the largest and growing populations worldwide of patients with T2DM (50), a metabolic disorder that is highly correlated with the development of TB and drug resistance (39). The statistical significance of any genomic epidemiological study of Mtb clinical samples in Mexico requires increasing the number of samples and the number of sites sampled, especially those states with no representative genomes.

With respect to the differences between the phenotypic and genotypic results related to drug-resistance, we found large discrepancies among our dataset of 133 *Mtb* samples. In Figure 5 we show the proportion of samples that agree and disagree in terms of antibiotic resistance by genotype (y-axis) and phenotype (panel A). Discrepancies between samples classified as susceptible according to a genotypic analysis but exhibiting a drug-resistant phenotype are particularly concerning in clinics (red arrow). According to a differential functional analysis between susceptible and resistant samples from these group, we found 26 genes containing possible variants, which had a fitness score greater than 1 (panel B). Among these genes, two genes of the PE and PPE family stand out (υ>2.14). The gene *ppe19* encodes for an uncharacterized protein while the *pe-pgrs17* gene encondes for a protein that has been reported as a functional protein during immune evasion by intervening in the antigen presentation process and, therefore, in the recognition of infected host cells and their elimination (51). Gene *ino1* plays a crucial role in the mycothiol biosynthesis in *Mtb*, which has been demonstrated as essential gene for growth and virulence in mice (52). This analysis suggests that the currently available databases of canonical variants associated with drug-resistance may be incomplete and that it is necessary to integrate new genomes from under-represented geographic areas in order to increase the accuracy of resistance prediction. On the other hand, discrepancies between phenotype and genotype could also be explained by the intrinsic limitations of the phenotypic methods, such as the use of single concentration tubes, or due to a selection of the fittest strains during expansion for DNA extraction. The mixture of susceptible and resistant strains in one single sample, known as heteroresistance (53), also has implications for the results of phenotypic DST (54). In addition to sequencing more representative genomes, the necessary technical controls must therefore be implemented in laboratory tests in order to more accurately detect antibiotic resistance.

**Figure 5.**
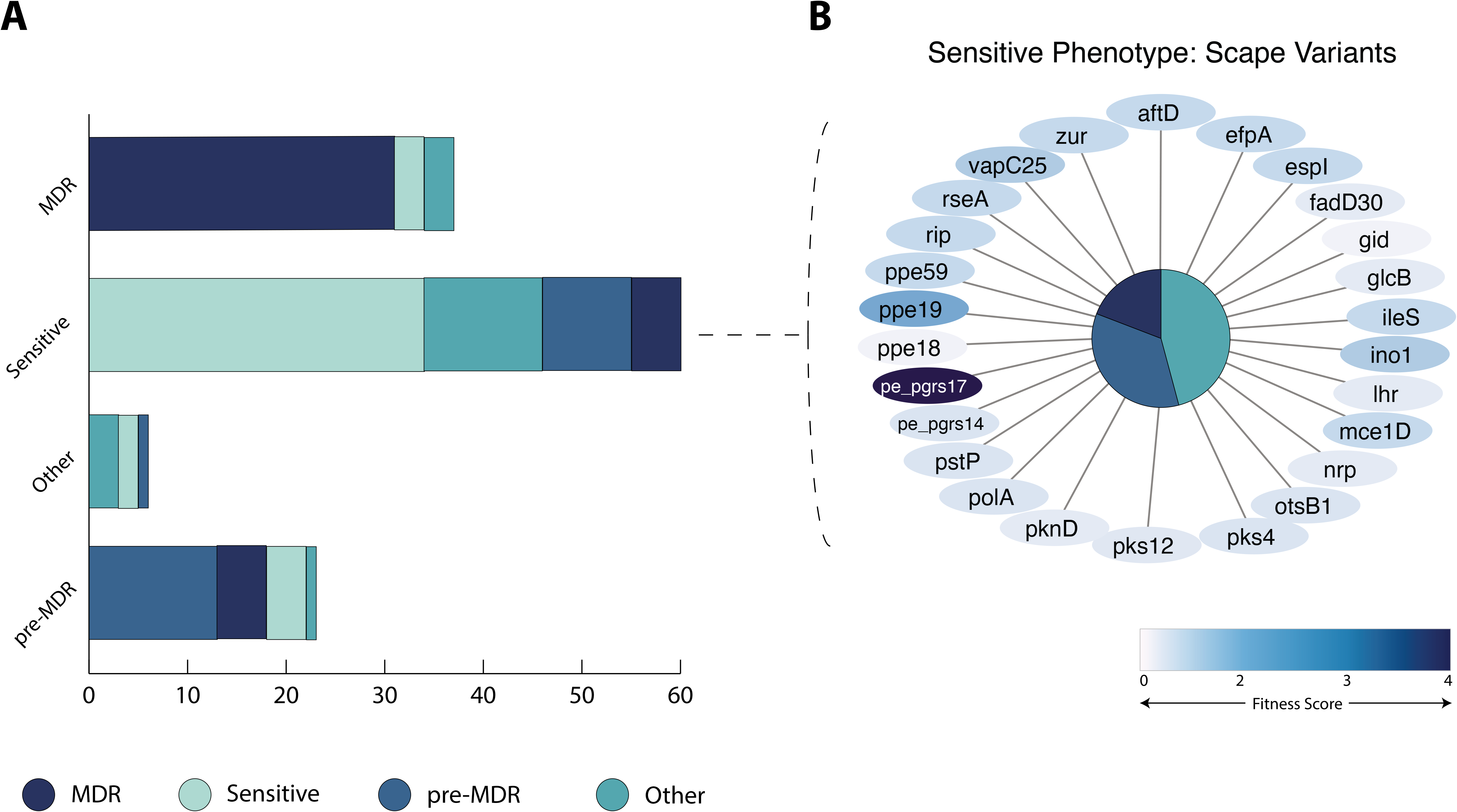
Representation of the contrasting results obtained from the phenotypic and genotypic drug-resistance classification for the 133 *Mtb* samples from Mexico. The main bar graph represents the genotypic classification of *Mtb* samples with respect to drug-resistance against first-line antibiotics. The secondary bar graphs show, for each class, the proportions of samples that agree or disagree based on their drug-resistance phenotypic class.

Aiming to understand the drug-resistance mechanisms that have evolved in *Mtb* strains isolated from Mexico, we conducted a functional analysis of all genes containing variable positions (non-synonymous changes) present only in strains with drug-resistant profiles. For this, we developed a bioinformatic tool (FuN-TB) to perform a comparative analysis of variants between any subset of samples in order to obtain a list of differentially mutated genes. The resulting list of genes can then be analyzed from a functional standpoint, thus providing a metabolic and cellular perspective of the acquired mutations. Newly sequenced genomes can therefore be easily integrated into the analysis if they comply with the basic genome sequencing requirements for *Mtb* strains (short-read sequencing technology, 65.6% GC, >98% reads mapped against the reference genome, coverage mean >25X). Sample subsets are created by selecting specific sample attributes or epidemiological metadata (e.g., DST phenotype, co-morbidity, geographic region, year of isolation, sex or age of the patient, type of treatment, etc.) of interest to the researcher. The FuN-TB tool is a standalone algorithm available to the scientific community interested in conducting comparative functional analysis of mutated genes among *Mtb* genomes, and with an understanding of the molecular basis of different attributes related to the samples.

In this study, we used the FuN-TB tool to analyze a set of 133 clinical samples of *Mtb* from Mexico in order to reveal the cellular and metabolic mechanisms (different from the global databases) that might be involved in drug-resistance adaptation in the region. According to the phenotype test, the first stage of this functional analysis aimed to identify the differentially mutated genes among five sub-sets of mono-resistant samples (H, R, E, S, or Z) according to the phenotype test. The exciting results produced by this mono-resistance functional analysis showed 31 mutated genes appearing simultaneously in the R and E mono-resistant samples, which remarkably shared the same mutation at the codon level. Among these, we found several genes involved in interaction with the host, response to host immune system, response to hypoxia and cell wall biosynthesis and maintenance (**Supplementary Table 5**). Human P-glycoprotein (P-gp) is an efflux pump expressed in lung epithelia which has been associated with a reduction in the intracellular accumulation of many drugs (55), including rifampicin and ethambutol, but not isoniazid (56). Recently, a rare polymorphism (2677G>A) was reported in the P-gp encoding gene from patients with pulmonary TB living in northeastern Mexico, which exhibited simultaneous resistance to rifampicin and ethambutol (57). These results sug According to the phenotype test, the first gest that human genetics in Mexico may influence the coordinated response of *Mtb* strains to rifampicin and ethambutol treatment. Further studies incorporating more significant number of *Mtb* isolates are required in order to confirm this association in the region.

Moreover, to identify the genes that contained frequent variable positions, we calculated a variation score (See **Methods**) for each gene of the mono-resistance analysis. The genes with the highest variation score were *hpt* and *etgB*, found in the E, R and S resistant subsets, as well as *ligC* and *pe35*, found in E, R and H resistant subsets of samples. The gene *hpt* is involved in the metabolism of the prodrug azathioprine (KEGG pathway, 58), an immunosuppressor acting as a tumor necrosis factor (TNF) antagonist. TNF is a cytokine involved in granuloma formation during *Mtb* infection (59). Treatments with TNF antagonists, such as azathioprine, increase the risk of progression of latent TB to active TB or re-activation of TB (60, 61). Interestingly, of our samples, four S resistant and one R resistant were relapsed TB cases. The gene *etgB* is part of the metabolic pathway of L-histidine, specifically involved in the biosynthesis of ergothioneine (EGT). EGT is a redox buffer derived from L-histidine that plays a key role during antibiotic resistance and virulence of *Mtb* by maintaining the redox and bioenergetic homeostasis of the cell under stress conditions (62, 63). The gene *ligC* encodes an ATP-dependent ligase, playing a role in BER and a backup role in NHEJ DNA repair systems (64, 65). Robust DNA repair systems are necessary for *Mtb* to overcome immune and antibiotic stresses (66). Most of the variants we found in the comparative analysis were not present in the recently published catalogue of drug-resistant variants (6) (**Supplementary Table 7**). These results highlight the importance of exploring beyond the canonical variants in the *Mtb* genomes. Considering the status of *Mtb* as one of the oldest pathogens of modern humanity, it is to be expected that its evolutionary mechanisms go beyond the target genes of anti-TB drugs.

### Data availability

The raw data from this study is available in the NCBI (National Centre for Biotechnology Information) database under ID.

## Acknowledgements

We wish to express our deepest gratitude to the individuals who agreed to participate in this study, and to the clinical personnel at all sample collection sites. This study was funded through several grants: Mexican National Council for Science and Technology (CONACYT) - Ciencia de Frontera – 319590 to LCC; postgrad scholarships to MPPM and RGAA from the CONACYT; a Seed Grant from Tecnológico de Monterrey to LCC, DDM and SA for genome sequencing. Thanks also go to StrainBiotech SAPI de CV for providing additional laboratory supplies.

## Authors’ contributions

MPPM, CDJE, EMJA and LCC designed all experiments and analysis. CGCY, NCJ, RGEJ, SHAR, VSF, ZCR, CDJE and EMJA extracted and collected genomic DNA and performed DST tests for all the clinical isolates. MPPM, DDCM, SA and LCC performed genome sequencing. MPPM, LREE, ZCR, CDJE, EMJA and LCC analyzed and interpreted all data. MLMF, LREE, CDJE and EMJA performed docking analysis. RGAA, MPPM and LCC designed and performed bioinformatic analysis. MPPM and LCC wrote the manuscript. All of the authors have read and critically reviewed the manuscript, and consent to its publication.

## Competing interests

The authors declare no conflict of interest.

## List of Supplementary Figures

**Supplementary Figure 1. Mutations outside of the GyrA Quinolone resistance determining region (QRDR) alter the polarity distribution at the fluoroquinolone binding site.** A) Comparison of the wild-type DNA Gyrase (Mtb H37Rv, green color) and the mutant (cyan color), showing that isolated amino acid mutations in GyrA do not lead to changes in protein structure. B) The topological polar surface area (TPSA) of DNA gyrase WT (red color) and strains with more than two mutations in GyrA: UVer054 (yellow color) and UVer059 (magenta color) changes the charge and polarity of the proteins. C) The predicted Vina binding energies decrease for the ligand docked for each protein isolated mutation. D) GyrA modelled with mutations in several positions from the clinical isolates, and the estimated binding energies for quinolone in the GyrA binding site.

## List of Supplementary Tables

**Supplementary Table 1. Epidemiological data from 133 *Mtb* genomes**

**Supplementary Table 2. Whole genome sequencing information from 133 *Mtb* genomes**

**Supplementary Table 3. Comparison of the sublineage designation for 53 *Mtb* samples**

**Supplementary Table 4. Discrepancies between the phenotype and genotype prediction of drug-resistance from 53 *Mtb* samples**

**Supplementary Table 5. Complete list of the mutated genes from the mono-resistant dataset analysis**

**Supplementary Table 6. Complete list of the mutated genes from the MDR dataset analysis**

**Supplementary Table 7. Comparison between variants found in our analysis and the WHO-catalogue of drug-resistant variants released in 2021**

